# Longitudinal analysis of innate immune system in infants with perinatal HIV infection until 18 months of age

**DOI:** 10.1101/2023.11.21.568007

**Authors:** Vinh Dinh, Lesley R. de Armas, Suresh Pallikkuth, Rajendra Pahwa, Stefano Rinaldi, Christine Dang, Alexander Kizhner, Nicola Cotugno, Paolo Palma, Nália Ismael, Paula Vaz, Maria Grazia Lain, Savita Pahwa

## Abstract

With the advent of antiretroviral therapy (ART), perinatal HIV infection is declining globally but prevalence in Sub-Saharan Africa is still greater than other nations. The relationship of HIV replication in early infancy and the developing immune system is not well understood. In this study, we investigated cellular components of the innate immune system including Natural Killer (NK) cells, monocytes, and Dendritic Cells (DC) in a cohort of HIV exposed infected (HEI) and age-matched HIV exposed uninfected (HEU) infants from Mozambique. Study entry was at the first visit after delivery at age 1-2 months for HIV diagnosis and initiation of ART. Phenotypic analysis by multi-parameter flow cytometry revealed an expansion of total NK cells and the dysfunctional, CD56^-^CD16^+^, NK cell subset; increased activation in monocytes and DC; and higher levels of inflammatory homing receptor CCR5 on circulating DC subsets in the HEI infants. NKG2A, an inhibitory receptor for NK cytolytic function, was reduced in HEI compared to HEU and positively correlated with pre-ART viral load (VL) while expression of CCR2, the inflammatory homing receptor, on NK was negatively correlated with VL. Other subsets exhibited positive correlations with VL including the frequency of intermediate monocytes amongst total monocytes. Longitudinal analysis of VL indicated suboptimal ART adherence in HEI. Regardless of level of viral suppression achieved, the frequencies of specific innate immune subsets in HEI were normalized to HEU by 18m. These data support the notion that in early life, NK cells play a role in virus control and should be explored for functional attributes that are effective against HIV at this time during development. Overall, our study provides high resolution overview of the innate immune system during perinatal HIV infection.

**Author Summary:** Vertical transmission of HIV has been reduced globally in recent years, however in utero exposure and acquisition of HIV continues to occur, especially in sub-Saharan Africa. Immediate ART initiation is recommended in infants diagnosed with HIV, but adherence is often suboptimal due to behavioral and sociological challenges. The impacts of perinatal HIV infection and ART on the developing immune system in infants are still unclear. Here, we evaluated a cohort of HIV exposed infected infants, and age-matched HIV exposed uninfected infants from Mozambique at pre-ART (age 1-2m) and post-ART longitudinally (up to 18m) specifically to compare the innate immune cellular components. We found that circulating innate immune cells including Natural Killer (NK) cells, monocytes, and Dendritic Cells (DC) exhibited altered distributions and more activated (inflammatory) phenotypes at pre-ART in infants with HIV suggesting the presence of a virus specific immune response. Despite suboptimal ART adherence in the cohort, differences in innate immune subsets between infected (suppressed and unsuppressed) and uninfected were not observed longitudinally pointing to normalized immune development despite HIV infection. Our study provides new insights into the early innate immune response during perinatal HIV.

## Introduction

Currently, 1.5 million children are living with HIV world-wide [1]. Even though perinatal transmission of HIV is preventable with maternal use of anti-retroviral therapy (ART), approximately 130,000 children with HIV (CWH) were born in 2022. In CWH, early ART initiation in infancy reduces the virus reservoir size with a delay in virus rebound in children if therapy is interrupted [2-5]. However, challenges in low resource countries often prevent optimal ART uptake and adherence [6]. How virus-host interaction impacts immune development in infants and children with HIV has not been fully elucidated.

The adaptive immune system in infancy is predominantly naïve, placing increased importance on the innate arm of the immune system which does not require priming [7]. A major component of the innate immune system is represented by NK cells which play a key role in anti-viral immunity. NK cells can be divided into 4 distinct functional and phenotypic subsets based on their expression of the lineage markers, CD56 and the activating receptor FcγRIII or CD16. The subsets are: 1) immature, cytokine-producing CD56^bright^CD16^-^ 2) intermediate, CD56^dim^CD16^-^ 3) mature, cytotoxic CD56^dim^CD16^+^ and 4) CD56^-^CD16^+^ cells, which expands in adults with chronic HIV infection [8-12]. IFNγ production by NK cells has been shown to have an inverse correlation with HIV reservoir size in virally suppressed adults [12, 13] and the CD56^dim^CD16^+^ NK cell subset expressing CD11b, CD161, and Siglec-7 is expanded in Elite controllers, pointing to a protective role of NK cell subsets in this subpopulation of people with HIV (PWH)[12].

Neonates have comparable, if not higher, numbers and frequencies of NK cells compared to adults and the distribution of major NK subsets is similar to adults [11, 14, 15]. NK cell subsets in infants also exhibit comparable expression of CD16, which mediates Antibody Dependent Cellular Cytotoxicity [15]. Although the expression of cytotoxic effector molecules granzyme B and perforin are also reported as being similar to adults, neonatal NK cells have comparatively less cytotoxic function [16, 17]. The decreased cytotoxicity has been attributed to increased expression of the inhibitory NK receptor NKG2A and a diminished ability to adhere to target cells due to insufficient expression of adhesion molecules including L-selectin and ICAM-1 [15, 18].

Other innate immune cells, namely Antigen Presenting Cells (APC) such as monocytes and DCs play a critical role to drive the adaptive immune response. Monocytes can be subdivided in to 3 major subsets, which are the Classical Monocytes (CM; CD14^+^CD16^-^) that are mainly primed for phagocytosis and innate sensing; Intermediate Monocytes (IM; CD14^+^CD16^+^) that express the highest levels of antigen and secrete pro-inflammatory cytokines, and the Non-Classical Monocytes (NCM; CD14^-^CD16^+^) that are geared towards complement and FcR mediated phagocytosis [19]. During early life, monocytes have reduced expression of adhesion molecules and pathogen recognition receptors (PRRs) such as the LPS sensing Toll-Like Receptor 4 (TLR4), with reduced release of TNFα and other pro-inflammatory cytokines and reduced activation of MyD88 and NF-kB phosphorylation when compared to adults [20-22]. In PWH, monocytes have been shown to play a critical role as inflammatory mediators in HIV-associated neurocognitive disorders and cardiovascular disease [23, 24]. In addition, monocytes/macrophages have also been implicated as a potential site of latent HIV reservoirs [23, 25]. Dendritic cells (DC) can be divided in to 2 distinct cell types; the conventional DC (cDC; CD11c^+^CD123^-^) which participate in antigen presentation and Plasmacytoid DC (pDC; CD11c^-^ CD123^+^) which play a role in the anti-viral response [26, 27]. The cDC subset can be subdivided in to 2 different subsets, cDC1 which have a high intrinsic capacity to cross-present antigens via MHC-I to activate CD8+ T cells and promote Th1 responses, and the cDC2, which express a wide range of sensing receptors such as TLRs, NOD-like receptor, and RIG-I-like receptors at levels similar to that of monocytes [28]. In comparison to adults, infant DCs have been shown to produce lower levels of pro-inflammatory cytokines, lower basal expression of MHCII and co-stimulatory markers (CD80/86) which are necessary for antigen presentation to T cells, and less responsiveness to IFN signaling [7, 27]. The cDC1 are the predominant subset in spleen and lymph nodes of infants while cDC2 are the predominant subset in these sites for adults [26]. The pDC in umbilical cord blood have a reduced polyfunctionality based on production of multiple cytokines [7, 26]. In HIV infection, DC have been reported to be a potential reservoir site and have been implicated in facilitating infection of CD4 T cell by trans-infection [29]. During acute HIV infection there is dysregulation of pDC anti-viral function and a reduction in their overall frequency, that are partially restored after initiation of ART [30, 31].

Immunologic investigations of CWH in infancy, especially on the innate immune system, have been limited but are crucial for designing cure or ART-free remission strategies. This is due to inherent difficulties in recruitment and sampling of blood and tissues from this population. The current study was conducted using peripheral blood samples collected from perinatally HIV exposed infected (HEI) and HIV exposed uninfected (HEU) infants in Maputo, Mozambique. The prospective study followed HEI and age matched HEU children longitudinally from 1 month until 2 years of age with examination of innate immune system over a period encompassing pre- and post-ART initiation.

## Methods

### Study participants

The cohort named TARA (Towards AIDS Remission Approaches) included 89 infants consisting of 46 HEU and 43 HEI who were enrolled between 2017-2019 in Maputo province, Mozambique [6]. Infants born to women with HIV were given post-natal HIV prophylaxis with Nevirapine in the first 6 weeks of age. All HIV exposed infants were tested for HIV diagnosis within 1-2 months of age on the Alere™ q HIV-1/2 molecular diagnostic platform, which is a simplified point of care HIV nucleic acid test. Infants were enrolled into the TARA study as HEI or HEU with parental consent. In HEI, triple drug ART was initiated with Zidovudine (AZT), Lamivudine (3TC) and Lopinavir/Ritonavir (LPV/rit) at diagnosis, median age 5.7 weeks (range: 4-10 wks). Follow-up visits were at 5, 10, and 18 months of age. HEU infants were tested for HIV at each visit up to 18 months of age to rule out of transmission by breastfeeding. All infants received routine vaccinations according to the National schedule including BCG at birth, DTaP-HepB-Hib-PCV10-OP at 2, 3 and 4 months of age and measles-rubella vaccine at 9 and 18 months of age. Plasma viral load and CD4 counts were obtained at study entry (**Table 1**) and throughout the follow-up period for HEI and HEU.

**Table 1:** HIV Exposed Uninfected (HEU) and HIV Exposed Infected (HEI) infants at Pre-ART initiation and follow up outcome.

|  | HEU | HEI |
| --- | --- | --- |
| N, enrolled | 46 | 43 |
| Completed follow-up | 34 | 35 |
| *Lost to follow-up* | 12 | 4 |
| *Died* | 0 | 4 |
| *Sero-converted* | 1 | 0 |
| Gender (M/F) | 17/17 | 13/22 |
| Age at Entry (wks) | 5.7 (4-10) | 5.7 (4-11) |
| VL log, n | N/A | 5.7+/-1.2 |
| CD4 c/mm3, n | N/A | 2211+/-909 |

### HIV virus load and absolute CD4 count measurement

Plasma HIV virus load (VL) and CD4 counts were determined in Maputo. Plasma VL was quantified using COBAS® AmpliPrep/COBAS® TaqMan® HIV-1 Test, version 2.0 (Roche Diagnostics, Germany) with a limit of detection at 20 copies/ml. Absolute CD4 counts were obtained in HEI using BD Multitest™ anti-CD3FITC/anti-CD8PE/anti-CD45PerCP/anti-CD4APC and TruCount tubes (Becton Dickinson, USA) and analyzed on a FACSCalibur flow cytometry (Becton Dickinson, USA).

### Sample collection and storage

Venous blood was processed within 2 hours in Maputo. Peripheral blood mononuclear cells (PBMCs) were isolated by Ficoll Hypaque fractionation and cryopreserved in liquid nitrogen. Plasma was aliquoted and stored at -80°C. Plasma and PBMC and were stored on site until shipment from Mozambique to the University of Miami on dry ice or in liquid nitrogen cryoshippers, respectively for further testing.

### Surface marker phenotyping of innate immune cells by flow cytometry

Cryopreserved PBMC were thawed and rested overnight in RPMI-1640 media containing L-glutamine, Pen-strep, and 10% Fetal Bovine Serum (FBS) at 37°C and 5% CO2. 1 to 1.5 million PBMC were treated with Human TruStain FcX (Biolegend) to block non-specific binding of fluorochrome-conjugated anti-human monoclonal antibodies to Fc receptors before being labeled with a 27-color panel of markers including the viability dye Live/Dead Blue (Invitrogen), following manufacturer’s protocol, and 26 fluorochrome-conjugated anti-human monoclonal antibodies (listed in **Table S1**) for 30 minutes at room temperature. All reagents were titrated on control PBMC for optimum concentration before usage. Labeled PBMC were then fixed in 1% paraformaldehyde (PFA) before acquisition on the 5 laser Cytek ® Aurora Flow Cytometer. Compensation beads (name and manufacturer) were used to create the single-color controls and prepared in the exact same manner as PBMC. Single-color bead controls and unstained cells were used for spectral unmixing and auto-fluorescence extraction in the SpectroFlo software (Cytek, version 2.2.0.4).

### Flow cytometry data analysis

Unmixed FCS files were exported for gating and analysis using FlowJo (BD, version 10.8.1). Manual gating of innate immune cell populations (NK, Monocytes, and Dendritic Cells) and surface marker expression was performed as follows and shown in **Fig. S1 and S2**. Total NK cells were identified from the CD45^+^CD3^-^CD20^-^CD14^-^CD123^-^HLA-DR^-^ population based on their CD56 and CD16 expression **(Fig. S1)**. NK cell subsets were further identified as CD56^bright^CD16, CD56^dim^CD16^-^, CD56^dim^CD16^+^, and CD56^-^CD16^+^. Antigen presenting cells were identified from the population of CD45^+^CD3^-^CD20^-^CD56^-^HLA-DR^+^ cells and classified based on CD14 and CD16 expression as total monocytes (CD14^+/dim^CD16^-/+^) or total Dendritic Cells (CD14^-^CD16^-^) **(Fig. S2)**. The 3 major monocytes subsets were gated from within the Total Monocyte gate and were classified as CM (CD14^+^CD16^-^), IM (CD14^+^CD16^+^) and NCM (CD14^dim^CD16^+^). DCs were classified based on CD11c and CD123 expression into CD11c^-^CD123^+^ pDCs and CD11c^+^CD123^-^ cDCs. The cDC were subdivided into cDC1 (CD1c^+^CD141^-^) and cDC2 (CD1c^-^CD141^+^). Surface markers related to activation (CD2, CD38, CD80, and CD86), inhibition (NKG2A, PD-L1, and TIGIT), homing/trafficking (CCR2 and CCR5), and lipid scavenging (CD36) were investigated for expression on each of the cell subsets listed above. Cell subsets with less than 50 cells were not analyzed for surface marker expression.

### NK cytotoxicity assay

PBMC were thawed and rested for 3 hours in RPMI-1640 media containing L-glutamine, Pen-strep, and 10% FBS at 37°C and 5% CO2. NK cells were purified by negative selection by using the EasySep™ Human NK cell Enrichment Kit (Stem Cell Technologies). 5×10^4^ NK were co-cultured with NK sensitive cell line, K562 (ATCC), or a T cell line infected with the SF2 dual tropic virus, HUT78/SF2 (NIH AIDS Reagent Program), that were labeled with CellTrace Far Red (Invitrogen) at an Effector to Target Ratio of 4:1 for 18 hours. Cultures were performed in the presence of fluorochrome conjugated anti-CD107a antibody to ‘capture’ degranulation events as described [32, 33]. Cells were harvested and stained with a viability dye Live/Dead Violet (Invitrogen), following manufacturer’s protocol, and 3 fluorochrome-conjugated anti-human monoclonal antibodies towards CD3, CD56, and CD16 for 30 minutes at room temperature. They were then resuspended in 1X PBS before acquisition on Sony SH800S Cell Sorter. The panel of antibodies are shown in **Table S2**

### Statistical analysis

Mann-Whitney U Test was utilized for cell frequency comparisons between study groups at all time points. Heatmaps were generated using Morpheus (https://software.broadinstitute.org/morpheus). Kruskal-Wallis test and Dunn’s multiple comparison test were used for a multiple comparison between the various groups based on viremic status at 10 and 18 months of age and frequency of subsets longitudinally. Spearman correlation was utilized to investigate the relationship between each phenotypic parameter and the pre-ART viral load. Statistical tests were performed using GraphPad Prism version 10.0.2 for Windows and results were considered statistically significant with a p value ≤0.05.

### Ethics statement

The study was approved by the National Bioethical Committee (IRB00002657, reference 102/CNBS/2016) and the Ministry of Health (MOH) in Mozambique and by the University of Miami institutional review board (IRB #20160493). All Infants’ legal caregivers consented for the participation in the study through written signed informed consent.

## Results

### Increased immune activation in NK cell subsets at pre-ART in HEI correlate with HIV virus load

To evaluate the impact of perinatal HIV infection on the NK cell compartment, we compared NK cell frequencies between HEI and HEU at study entry (pre-ART, 1-2 months of age). At this timepoint HEI had viral loads ranging from 695 to 1 x 10^7^ copies/ml (**Fig. 1A)**. Peripheral blood NK cells and their subsets in blood were identified based on the expression of CD56 and CD16 (**Fig. 1B**). HEI infants had a higher frequency of total NK cells out of live CD45+ lymphocytes compared to HEU (median: 11.4% vs 9.3%) and were enriched for the CD56^-^CD16^+^ NK cell subset (median: 10.3% vs 6.6%), while the frequencies of CD56^bright^CD16^-^ (median: 4.2% vs 6.4%) and CD56^dim^CD16^-^ (median: 8.6% vs 12.2%) subsets were lower in HEI compared to the HEU. As in adults, the CD56^dim^CD16^+^ NK subset was predominant, representing a median of 67% and 65% of all NK in both HEI and HEU, respectively (**Fig. 1C**).

**Figure 1:**
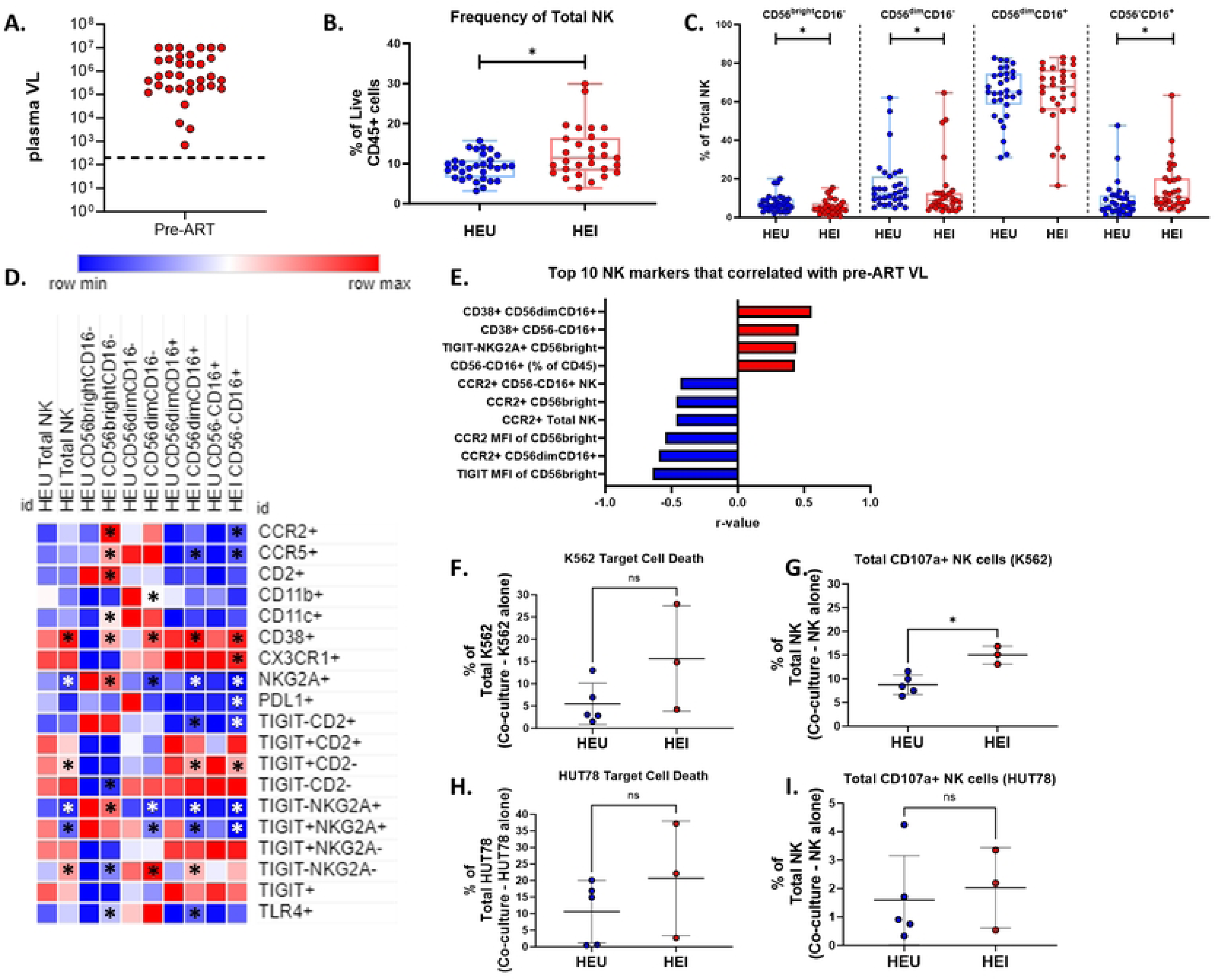
HEI infants have enriched NK cell subset frequencies with immune activation at pre-ART compared to age matched HEU that correlated with plasma viral load. A) Pre-ART VL of HIV-exposed infected infant (N=31) B) Frequency of NK cells as a percentage of live CD45+ cells in HIV-exposed uninfected (HEU, blue circles, N=31) and HIV-exposed infected (HEI, red circles, N=29). C) Frequencies of NK cell subsets based on CD56 and CD16 expression as a percentage of Total NK cells for both HEU and HEI. D) Heatmap generated using average frequency of each marker for each respective NK subset in HEU and HEI without any normalization. The color gradient of the heatmap represents the average frequency of a marker between subsets with blue being the lowest frequency and red being the highest frequency. T-test comparisons were done between study groups for each marker within a row. Any significant difference for a specific marker of each subset between study groups was marked with an asterisk on the heatmap. E) Top 10 most significant NK cell parameters that correlated with Pre-ART VL based on lowest p-value from the Spearman Correlation and r-values of each of these correlations is plotted across the x-axis. F-G) Target Cell Death and Total CD107a of NK cells when co-cultured with K562 target cells. H-I) Target Cell Death and Total CD107a of NK cells when co-cultured with HUT78 target cells.

We examined phenotypic markers of activation, inhibition, and trafficking on the NK cell subsets. A heatmap was generated to visualize the difference in mean frequency of these markers across the subsets between HEU and HEI. Comparisons between HEU and HEI for specific subsets and markers are depicted in **Fig. 1D and Table S3**. In HEI as compared to HEU, total NK cells and all subsets displayed higher frequency of the immune activation marker CD38 and lower frequency of the inhibitory receptor NKG2A. Differences were also noted in trafficking receptors CCR2 and CCR5; in HEI, higher frequencies of CCR2 and CCR5 expressing cells were observed on multiple subsets (**Fig. 1D and Table S3)**. Decreased frequency of NK cells that were either TIGIT+CD2-, TIGIT-NKG2A+, or TIGIT+NKG2A+ were observed in HEI when compared to HEU (**Fig. 1D and Table S3)**.

Next, we investigated whether the size (frequency) of any NK cell subset or parameter was associated with pre-ART plasma VL. Out of 176 NK related markers, both frequency and Median Fluorescence Intensity (MFI) data tested for univariate correlations with plasma VL, 22 had significant associations **(Fig. 1E and Table S4)**. Included among the top 10 NK parameters that correlated with VL were negative correlations of TIGIT MFI on CD56^bright^CD16^-^ and CCR2 frequency and MFI on Total NK, CD56^bright^CD16^-^, and CD56^dim^CD16^+.^subsets. Positive correlations were observed for frequency of CD56^-^CD16^+^ out of Live CD45+ cells, as well as frequency of CD38 on CD56^dim^CD16^+^ and CD56^-^CD16^+^ and of TIGIT-NKG2A+ on CD56^bright^CD16^-^ cells (**Fig. 1E**). In addition to the top 10 markers, CD38 MFI on Total NK and CD56^dim^CD16^+^ and NKG2A MFI on CD56^bright^CD16^-^ showed positive correlations, while MFI of CCR5 on Total NK and CD56^-^CD16^+^ and CCR2 MFI on CD56^dim^CD16^-^ cells were negatively correlated with Pre-ART VL (**Table S4**). Given the activated phenotype and associations with plasma VL, we investigated the cytotoxic capacity of NK cells and their ability to degranulate when co-cultured with either the MHC-I deficient K562 cell line or the HIV-infected HUT78/SF2 cell line in a portion of HEU and HEI participants. NK from HEI and HEU had a similar cytotoxic capacity, though the frequency of degranulating cells in response to K562 was higher in HEI. (**Fig. 1F-I**).

Taken together, the results show that NK cells from HEI infants at pre-ART initiation have skewed differentiation pattern and an altered phenotypic profile compared to HEU with the expansion of CD56^-^CD16^+^ being positively associated with ongoing viral replication. CD38 and CCR2 expression were higher in HEI compared to HEU, but these two markers exhibited different associations, CD38 being positively correlated and CCR2 being negatively correlated with pre-ART VL suggesting viral replication is associated with the activation of NK cells and increased trafficking by CCR2+ NK cells may be associated with viral control.

### NK cell compartment showed a transition to a profile similar to HEU during the first 18 months of life

Next, we assessed how NK cell development was impacted by the presence of HIV and ART initiation. Participants in the TARA cohort showed suboptimal adherence to ART therapy with a majority not adhering as reflected in lack of consistent virus suppression without evidence of resistance mutations [6]. We investigated how NK parameters that were altered in HEI at pre-ART initiation changed over time in HEI and HEU. At 10 months of age, a subset (N=11) of the HEI achieved viral suppression, defined as having a plasma viral load under 200 copies/mL, while the rest remained viremic (**Fig. 2A**). At this timepoint, the frequency of total NK cells from HEI, both viremic and aviremic, were similar to each other and to HEU (**Fig. 2B**). This pattern extended to the major subsets except the CD56^-^CD16^+^ subset in which viremic HEI maintained an elevated frequency relative to age-matched HEU, while the aviremic became similar to the HEU (**Fig. 2C**).

**Figure 2:**
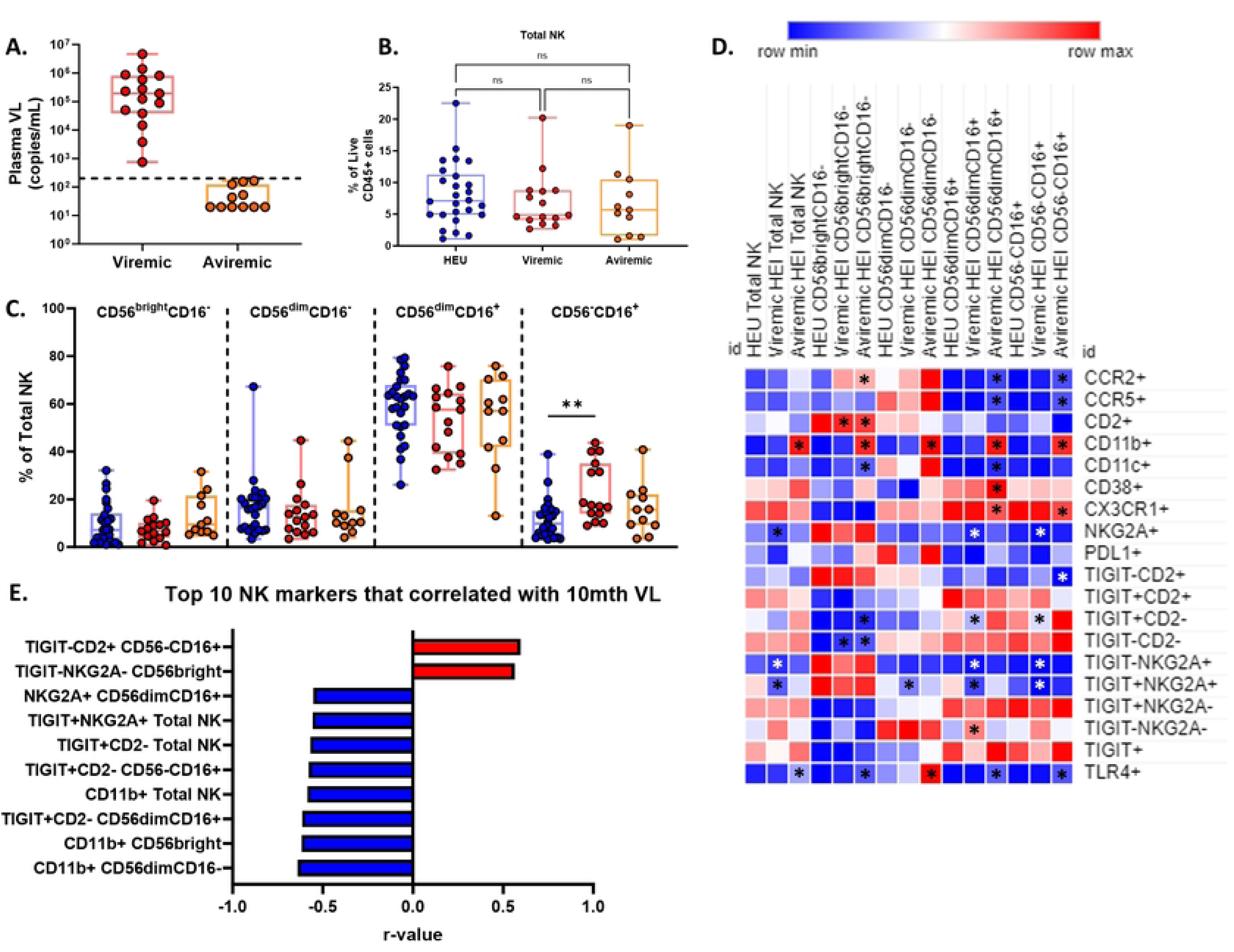
HEI infants maintain an enriched CD56-CD16+ subset and elevated immune activation at 10 months of age compared to age matched HEU that correlated with plasma viral load. A) HEI infant were split in to 2 groups at 10 months of age based on their 10-month VL. Viremic infants were defined by having a viral load above 200 copies/mL (N=15) and Aviremic infants were defined by having a viral load below 200 copies/mL (N=11). B) Frequency of NK cells as a percentage of live CD45+ cells in HEU (Blue circles, N=26), Viremic HEI (Red circles, N=15), and Aviremic HEI (Orange circles, N=11). C) Frequencies of NK cell subsets based on CD56 and CD16 expression as a percentage of Total NK cells for HEU, Viremic HEI, and Aviremic HEI. D) Heatmap generated using average frequency of each marker for each respective NK subset in HEU and HEI without any normalization. The color gradient of the heatmap represents the average frequency of a marker between subsets with blue being the lowest frequency and red being the highest frequency. T-test comparisons were done between study groups for each marker within a row. Any significant difference for a specific marker of each subset between HEU and either Viremic or Aviremic HEI was marked with an asterisk on the heatmap. E) Top 10 most significant NK cell parameters that correlated with 10-mth VL based on lowest p-value from the Spearman Correlation and r-values of each of these correlations is plotted across the x-axis.

Analysis of NK cells expressing markers of activation, inhibition, and trafficking in HEU and viremic or aviremic HEI at 10 months of age revealed a reversal of some of the differences exhibited at pre-ART initiation, such as the elevated frequency of CD38+ NK cells at Pre-ART initiation in all subsets observed in HEI was only elevated in CD56^dim^CD16^+^ NK cells of aviremic individuals (**Fig. 2D and Table S5**). Higher frequency of cells positive for trafficking markers, CCR2 and CCR5, in a few subsets of NK cells remained at this timepoint such as the CD56^-^ CD16^+^ NK cells in aviremic HEI and decreased frequency of cells positive for NKG2A was still maintained in the CD56^dim^CD16^+^ and CD56^-^CD16^+^ NK cells of Viremic HEI (**Fig. 2D**). Overall, by 10 months of age NK cells of HEI, both viremic and aviremic, maintained an elevated activation profile compared to HEU but not to the same extent as Pre-ART initiation. The markers associated with plasma VL at this time included inhibitory receptors, such as NKG2A and TIGIT, which negatively correlated with VL, and activation markers CD38 and CD2 that were positively correlated with VL (**Fig. 2E; Table S6**).

To investigate the effect of HIV and viremic status on development of the innate immune system during the first 2 years of life, the HEI were split in to 3 groups based on when, and if, they achieved viral suppression. HEI that never achieved a viral load below 200 copies/mL were classified as viremic (N=13), HEI that achieved a viral load under 200 copies/mL after 12 months of age and maintained it or showed occasional suppression during the first 2 years of life were classified as Partial/Late suppressors (P/L; N=10), and HEI that achieved viral suppression before 12 months with no more than one rebound event were classified as Early Suppressors (N=10). Viral load trajectories for the 3 HEI groups can be seen in **Fig. S3**. Longitudinally, P/L showed a decrease of Total NK cells with increasing age, and HEU showed a transient increase of CD56^-^ CD16^+^ during the first 5 months of age, while other subsets in all 4 participant groups were not associated with age (**Fig. 3A-E**). In cross-sectional analysis at 18 months of age, only 6 NK populations from either the viremic or P/L showed significant differences compared with HEU (**Fig. 3F; Table S7**). Viremic HEI exhibited a decrease of NKG2A+ CD56^dim^CD16^+^, TIGIT+NKG2A+ Total NK and CD56dimCD16+ NK cells, and CX3CR1+ CD56^bright^CD16^-^ NK cells (**Fig. 3F**). P/L showed a significant increase in frequencies of CCR5 expressing CD56^bright^CD16^-^ and CD56^-^CD16^+^ NK cells when compared to HEU. Overall, regardless of VL trajectories in HEI there was a reversal of phenotypes that were observed at pre-ART initiation towards a profile similar to HEU by 18 months of age.

**Figure 3:**
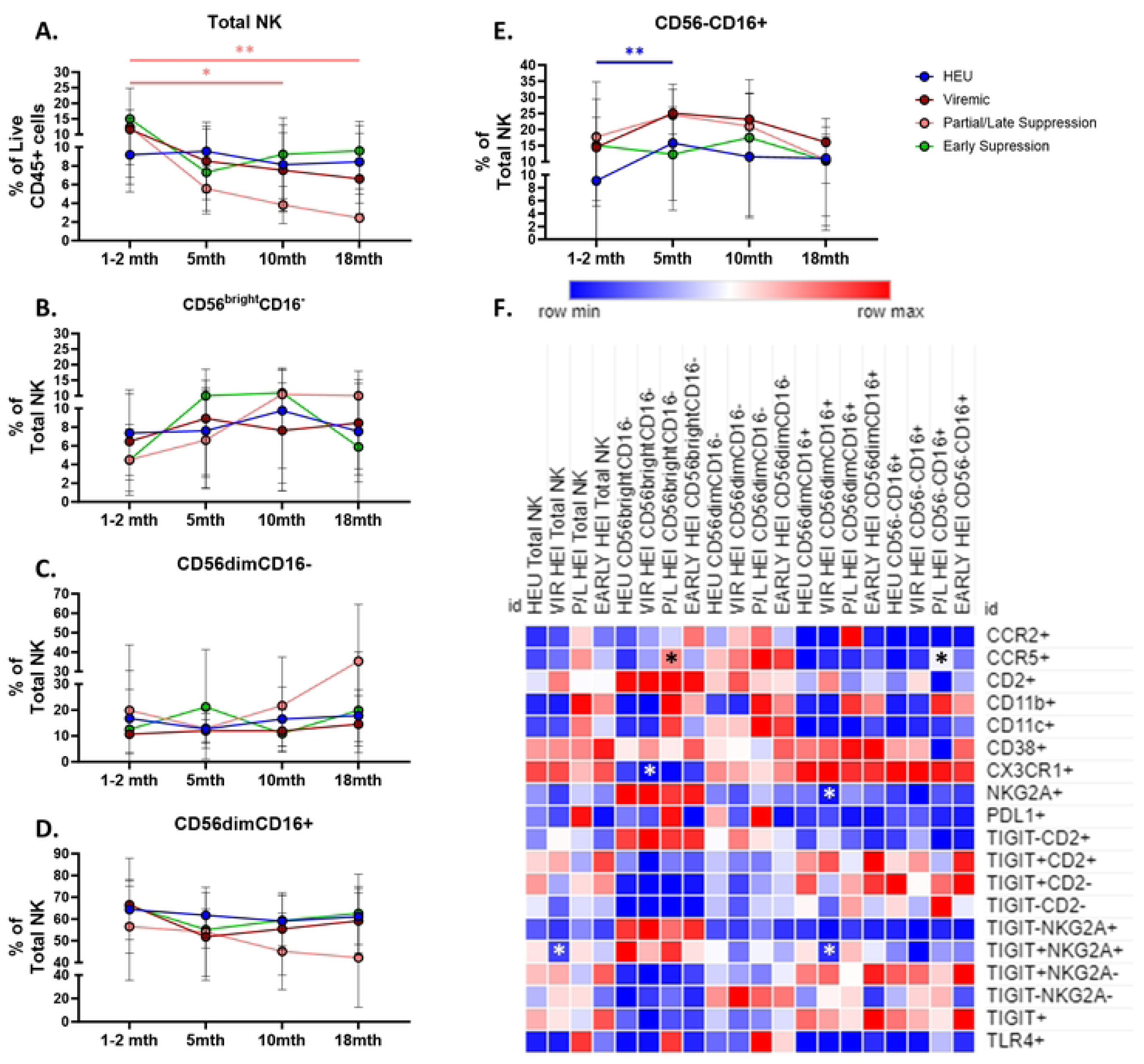
Regardless of viral trajectory, HEI show minimal differences in subset distribution and phenotype at 18 months of age compared to age matched HEU. HEI infants were split by their plasma viral load trajectories as shown in **Figure S3**. A) Frequency of NK cells as a percentage of live CD45+ cells in HEU (Blue circles, N=26), Viremic (Dark Red circles, N=13), Partial/Late Suppressors (Pink Circles, N=10) and Early Suppressors (Green circles, N=10). B-E) Frequencies of NK cell subsets based on CD56 and CD16 expression as a percentage of Total NK cells for HEU, Viremic, Partial/Late suppressors and Early Suppressors. D) Heatmap generated using average frequency of each marker for each respective NK subset in HEU and HEI without any normalization. The color gradient of the heatmap represents the average frequency of a marker between subsets with blue being the lowest frequency and red being the highest frequency. T-test Comparison were done between study groups for each marker within a row. Any significant difference for a specific marker of each subset between HEU and any of the 3 HEI groups was marked with an asterisk on the heatmap.

### Monocyte subsets exhibit increased activation and correlate with pre-ART VL

Monocytes play an integral role in the innate immune response and have been shown to be a potential target for HIV infection, therefore we investigated the distribution of monocyte subsets between HEI and HEU and performed correlation analysis of subsets with pre-ART VL among HEI. Total Monocytes as well as the 3 major monocyte subsets (CM, IM, and NCM) were similar in frequency in HEI and HEU at pre-ART initiation (**Fig. 4A-B**). Utilizing the same approach as NK compartment, we investigated the differences between HEI and HEU in the frequency of monocytes expressing activation, inhibitory, trafficking, and scavenging receptors at pre-ART initiation (**Fig. 4C and Table S8**). Total and NC monocytes had elevated expression of the immune activation marker CD38 in HEI (**Fig. 4C**). In addition, HEI had a higher frequency of the lipid scavenging marker, CD36 on TM and NCM compared to HEU. The frequency of monocytes expressing high affinity scavenger receptor, CD163 on TM, CM, and IM was lower in HEI compared to HEU which could be explained by shedding of the surface marker following activation. Expression of CCR2 on TM, CM, and IM was reduced in HEI however CCR5, the HIV coreceptor and homing molecule for sites of inflammation did not differ between HEI and HEU among monocyte subsets.

**Figure 4:**
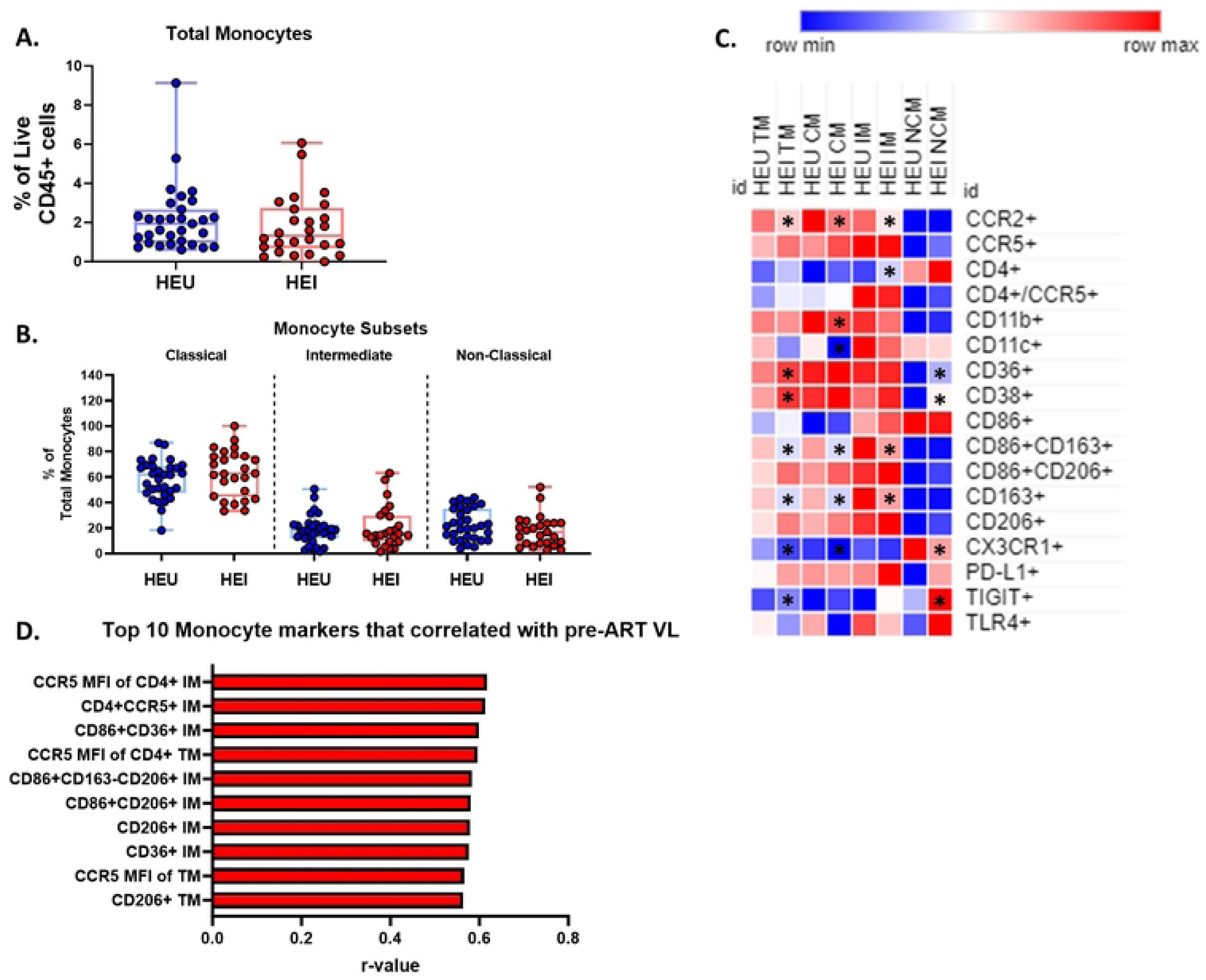
Monocytes show increased activation when compared to age matched HEU at Pre-ART initiation and Intermediate monocytes correlated with pre-ART VL. A-B) Total Monocytes as a frequency of Live CD45+ cells and frequency of Monocyte subsets as a percentage of Total Monocytes in HIV-exposed uninfected (HEU, blue circles, N=31) and HIV-exposed infected (HEI, red circles, N=29). C) Heatmap generated using average frequency of each marker for each respective Monocyte subset in HEU and HEI without any normalization. The color gradient of the heatmap represents the average frequency of a marker between subsets with blue being the lowest frequency and red being the highest frequency. T-test Comparison were done between study groups for each marker within a row. Any significant difference for a specific marker of each subset between study groups was marked with an asterisk on the heatmap. D) Top 10 Monocyte parameters that correlated with Pre-ART VL based on lowest p-value from the Spearman Correlation and r-values of each of these correlations is plotted across the x-axis.

Out of 280 monocyte related parameters, both frequency and MFI, tested for univariate correlations with plasma VL, 33 had significant associations. The top 10 correlations, based on p-values, with pre-ART VL included IM expressing either CCR5, CD4, CD36, CD86, CD163, and CD206 and these were positively correlated with VL. (**Fig. 4D**). Outside of the top 10 markers based on p-values, IM, as well as CM and NCM, expressing CD38, CD11b, CD36, also showed positive correlations with VL (**Table S9**). Expression of CCR5 (frequency and MFI) on TM, CM, and IM and mannose receptor CD206 on all subsets of monocytes showed significant positive correlation with plasma VL, despite no difference in expression of either molecule relative to HEU at pre-ART initiation. (**Fig. 4D and Table S9**). The data suggests that monocyte distribution is not significantly impacted by HIV in infants, but multiple IM phenotypes were positively associated with VL suggesting that this subset is influenced by HIV viral replication.

### Longitudinal analysis of monocyte compartment

To investigate the effect of ART initiation on monocyte development in infants, HEI were split into Viremic and Aviremic groups based on VL at 10 months of age (**Fig. 2A**). The frequency of Total Monocytes and distribution of the 3 major subsets (CM, IM, and NCM) were similar between the HEU, viremic HEI, and aviremic HEI (**Fig. 5A-B**). Expression of surface markers at this timepoint revealed primarily differences between HEU and aviremic HEI, rather than HEU and viremic HEI (**Fig. 5C; Table S10**). The monocyte phenotypes observed at pre-ART initiation in HEI were mostly restored to match HEU levels at 10 mo such as elevated expression of CD38 and CD86, and decreased frequency of CCR2+ monocytes (**Fig. 4C and 5C**).

**Figure 5:**
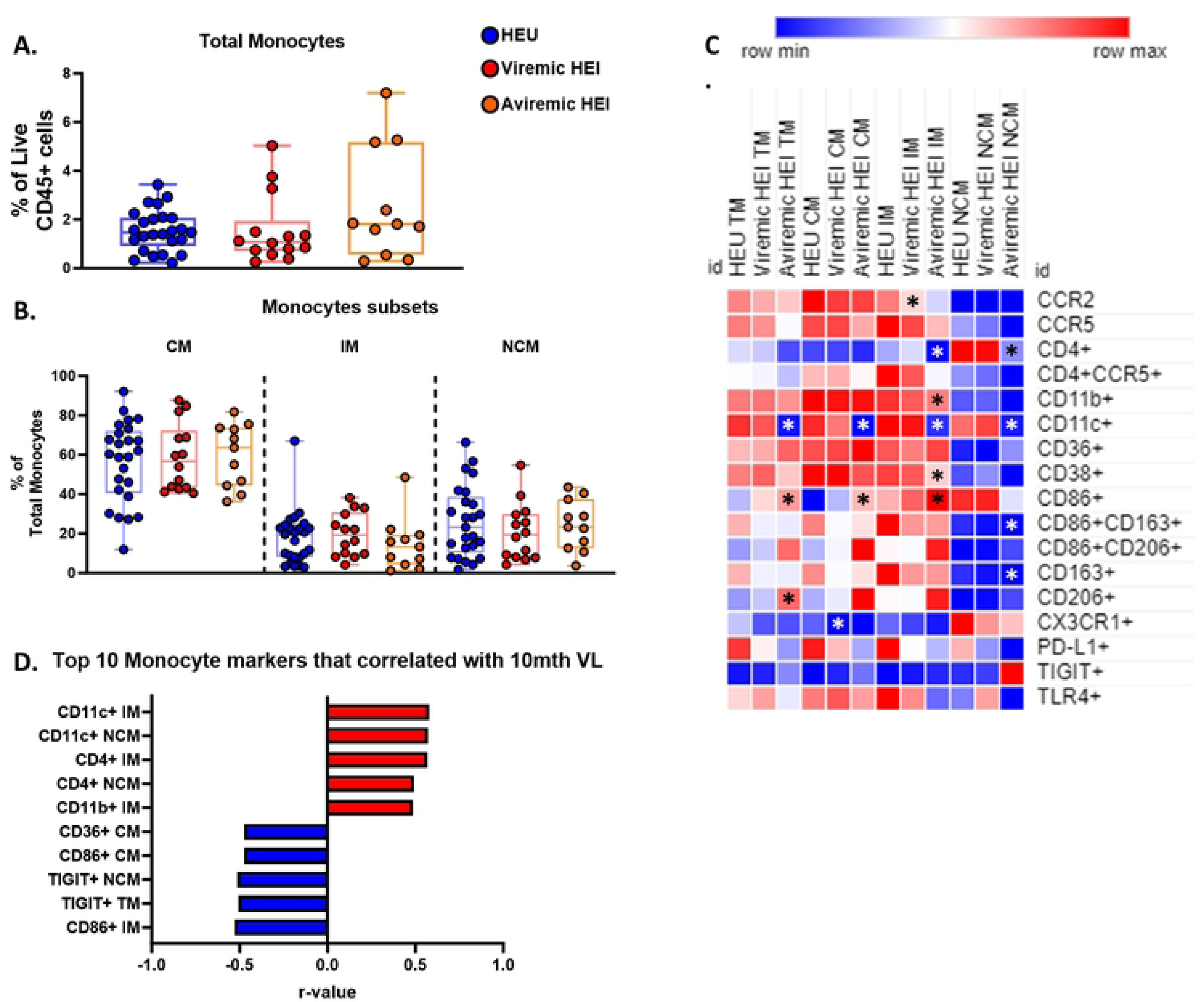
Aviremic HEI at 10 months of age show phenotypic differences from HEU and Intermediate monocytes correlate with plasma VL. HEI infant were split in to 2 groups at 10 months of age based on their 10-month VL. A-B) Total Monocytes as a frequency of Total CD45+ cells and frequency of Monocyte subsets as a percentage of Total Monocytes in HIV-exposed uninfected (HEU, blue circles, N=25), Viremic HIV-exposed infected (Viremic HEI, red circles, N=14), and Aviremic HIV-exposed infected (Aviremic HEI, Orange circles, N=11). C) Heatmap generated using average frequency of each marker for each respective Monocyte subset in HEU and HEI without any normalization. The color gradient of the heatmap represents the average frequency of a marker between subsets with blue being the lowest frequency and red being the highest frequency. T-test Comparison were done between study groups for each marker within a row. Any significant difference for a specific marker of each subset between HEU and either Viremic or Aviremic HEI was marked with an asterisk on the heatmap. D) Top 10 most significant Monocyte parameters that correlated with 10-month VL based on lowest p-value that from the Spearman Correlation and r-values of each of these correlations is plotted across the x-axis.

Correlation analysis with VL at this timepoint revealed positive correlation between CD38 expression and negative correlation of CD86 expression (**Fig. 5C; Table S11**). In addition, IM and NCM that expressed CD4 also positively correlated with plasma VL (**Fig. 5D; Table S10**). Interestingly, aviremic HEI had a decreased frequency of cells expressing the integrin, CD11c across all subsets. CD11c was also found to positively associated with VL when including all HEI infants. (**Fig. 5C; Table S11**).

Viremic, Partial/Late, and Early Suppressing HEI were assessed for longitudinal changes in the monocyte subset distribution and compared to the longitudinal changes observed in HEU. No significant differences were observed between the study groups, but IM showed a significant decline in frequency during the first 18 months of life, as well as NCM from 10 to 18 months of age (**Fig. 6A-D**). Cross-sectional analysis at 18 months of age demonstrated that monocyte phenotype was not impacted in HEI regardless of viral suppression history or status at this age (**Fig. 6E; Table S12**).

**Figure 6:**
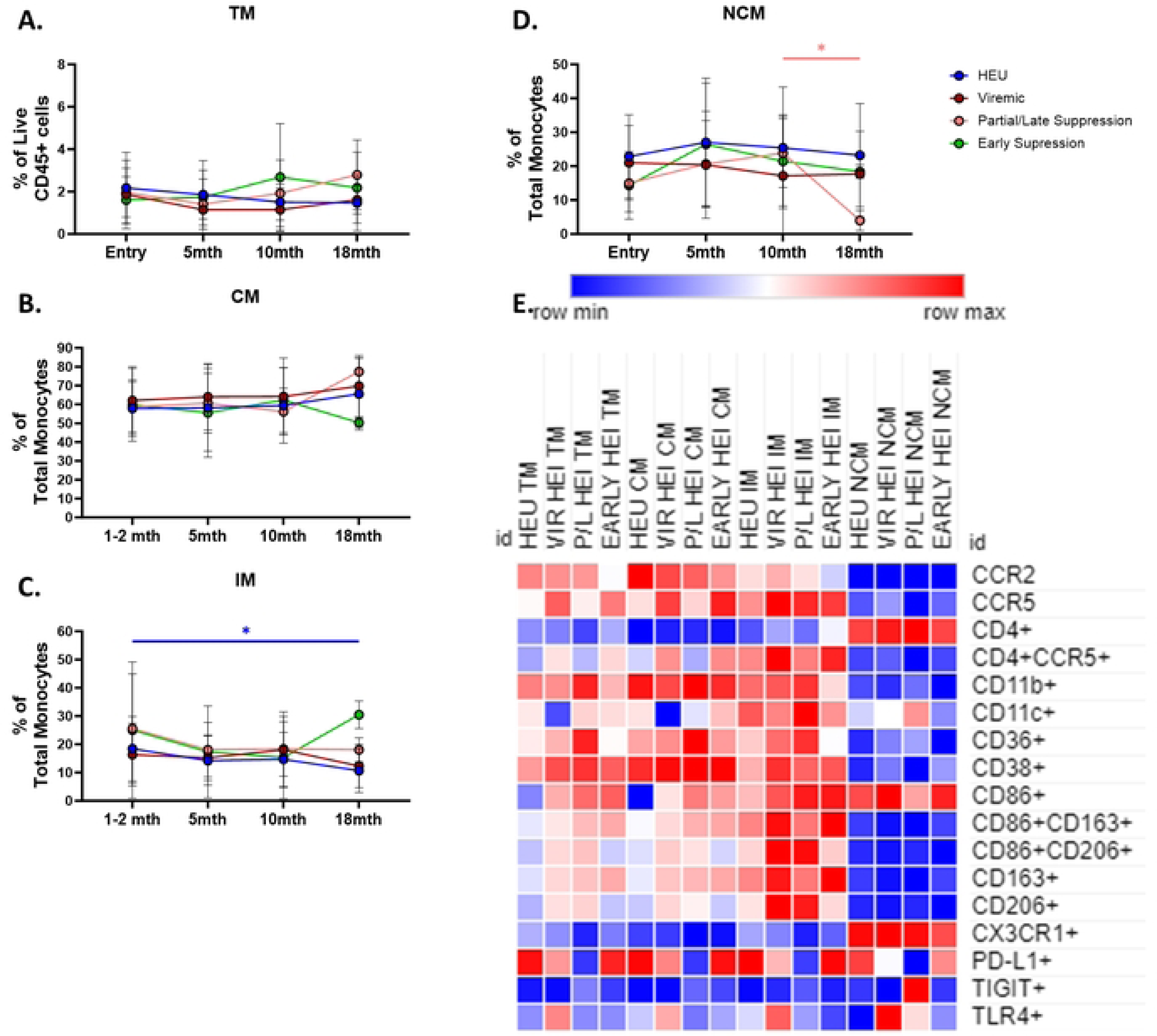
Regardless of viral trajectory, HEI show minimal differences in Monocyte subset distribution and phenotype at 18 months of age compared to age matched HEU. HEI infants were split by their plasma viral load trajectories as shown in **Figure S3**. A-D) Total Monocytes as a frequency of Live CD45+ cells and frequency of Monocyte subsets as a percentage of Total Monocytes based on expression of CD14 and CD16 in HEU (Blue circles, N=26), Viremic (Dark Red circles, N=13), Partial/Late Suppressors (Pink Circles, N=10) and Early Suppressors (Green circles, N=10) plotted against age in months on the x-axis. D) Heatmap generated using average frequency of each marker for each respective Monocyte subset in HEU and HEI without any normalization. The color gradient of the heatmap represents the average frequency of a marker between subsets with blue being the lowest frequency and red being the highest frequency. T-test Comparison were done between study groups for each marker within a row. Any significant difference for a specific marker of each subset between HEU and any of the 3 HEI groups was marked with an asterisk on the heatmap.

### Dendritic cells of HEI at Pre-ART initiation exhibit an altered phenotype compared to HEU

DC are important innate immune cells that act as a bridge between the Innate and adaptive immune system and like Monocytes can be a target for HIV infection, therefore next, we compared conventional DC (cDC) and plasmacytoid DC (pDC) between HEI and HEU and determined associations with pre-ART VL in HEI. There was no significant difference in the frequency of Total DCs and pDCs between HEU and HEI, while the frequency of cDCs were lower in HEI, and the frequency of cDC2, of total cDC, were higher (**Fig. 7A-C**). Markers of activation, inhibition and trafficking were evaluated on DC subsets and showed that similar to monocytes, expression of CCR2 was reduced in Total DC, pDC and cDC in HEI (**Fig.7D and Table S13**). Among additional markers on DC, lower frequencies were noted in HEI for the LPS recognition receptor TLR4 in Total DC and subsets (**Fig. 7D**), integrin CD11b, and co-stimulatory marker CD86 on Total DC and cDC (**Fig. 7D**). Total cDC, including those expressing CD4, exhibited heightened expression of CCR5 compared to HEU. (**Fig. 7D)**. Overall, expression of CD38 and CD86 on DC were significantly decreased in HEI compared to HEU.

**Figure 7:**
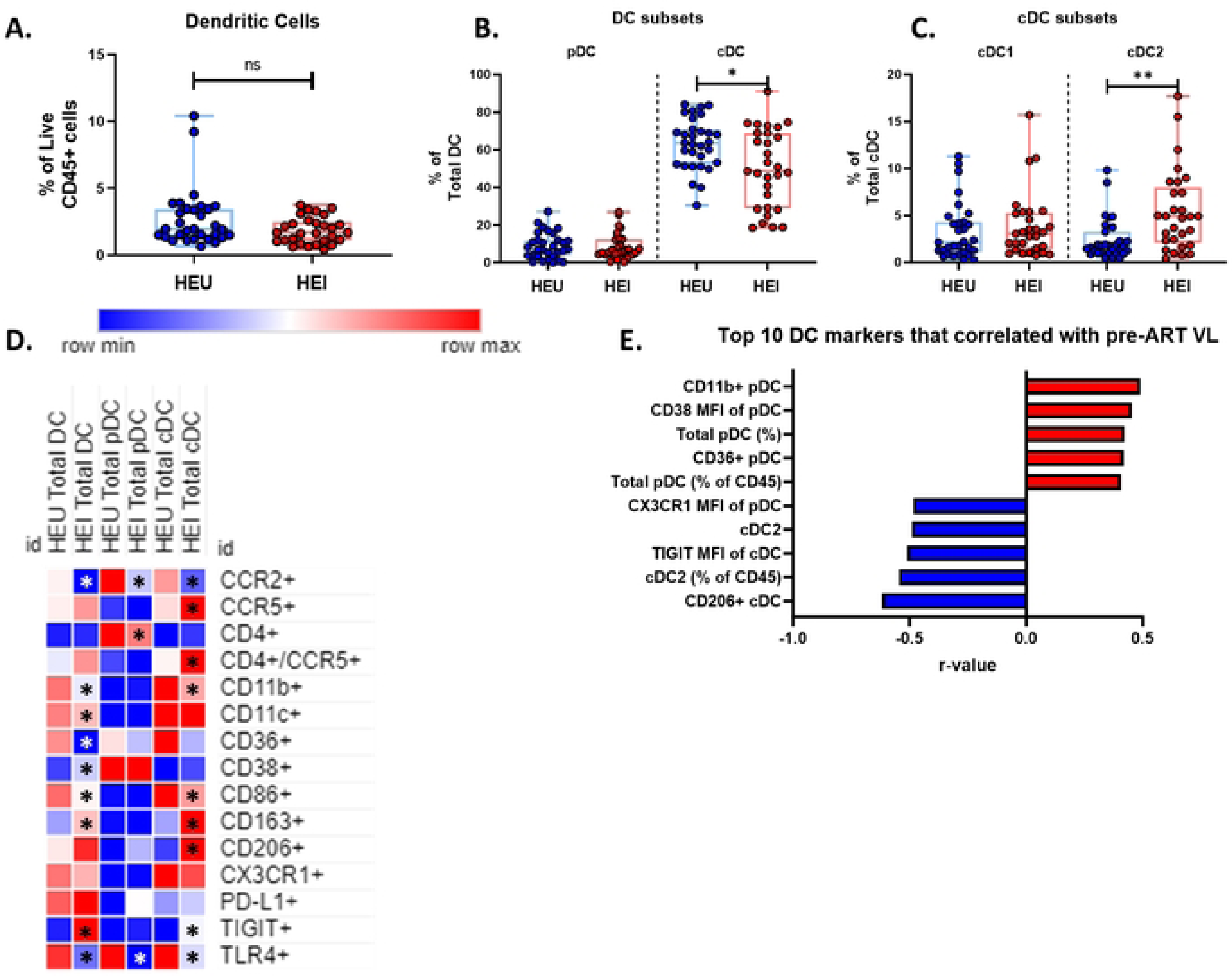
Dendritic cells of HEI at pre-ART initiation show an altered phenotype compared to HEU and activation related markers correlate with plasma VL. A-C) Frequency of Dendritic cells as a frequency of Live CD45+ cells, pDC and cDC as a percentage of Total Dendritic Cells, and cDC2 and cDC1 as a frequency of total cDC in HIV-exposed uninfected (HEU, blue circles, N=31) and HIV-exposed infected (HEI, red circles, N=29). D) Heatmap generated using average frequency of each marker for each respective Dendritic Cell subset in HEU and HEI without any normalization. The color gradient of the heatmap represents the average frequency of a marker between subsets with blue being the lowest frequency and red being the highest frequency. T-test Comparison were done between study groups for each marker within a row. Any significant difference for a specific marker of each subset between study groups was marked with an asterisk on the heatmap. E) Top 10 most significant Dendritic Cell parameters that correlated with Pre-ART VL based on lowest p-value that from the Spearman Correlation and r-values of each of these correlations is plotted across the x-axis.

The top 10 most significant associations of DC with pre-ART VL, out of 115 DC parameters, (**Fig. 7E; Table S14**) included negative correlations with CD206+ cDC and cDC2 as well as CX3CR1 MFI on pDC. Positive correlations were noted for pDC, both as a frequency of Total DC and total CD45+ cells, and expression of CD11b, CD36, and CD38 on this subset. Additional correlations outside of the top 10 markers, activation marker CD38 and CD86 on Total DC and cDC showed positive correlation with pre-ART VL and inhibitory receptor TIGIT on Total DC and pDC showed a negative correlation. Taken together, as compared to HEU, the HEI manifested an altered distribution of DC subsets with increased activation that was associated with the Pre-ART VL.

### Effect of ART initiation on DC phenotypes

We analyzed longitudinal data on the DC compartment to investigate the impact of HIV infection and ART initiation on immune development. At 10 months of age, no significant differences were observed in the frequency of DC subsets in Viremic and Aviremic HEI compared to HEU (**Fig.8A-C**). Surface marker analysis within DC subsets showed that Aviremic HEI, rather than Viremic HEI had a distinct profile relative to HEU (**Fig. 8D; Table S15**). Within pDC there was elevated expression of the integrin CD11b, co-stimulatory marker CD86, and scavenging receptors CD163 and CD206, while cDCs had elevated CCR5 expression on both total cDC and CD4+ cDC (**Fig. 8D; Table S15**).

**Figure 8:**
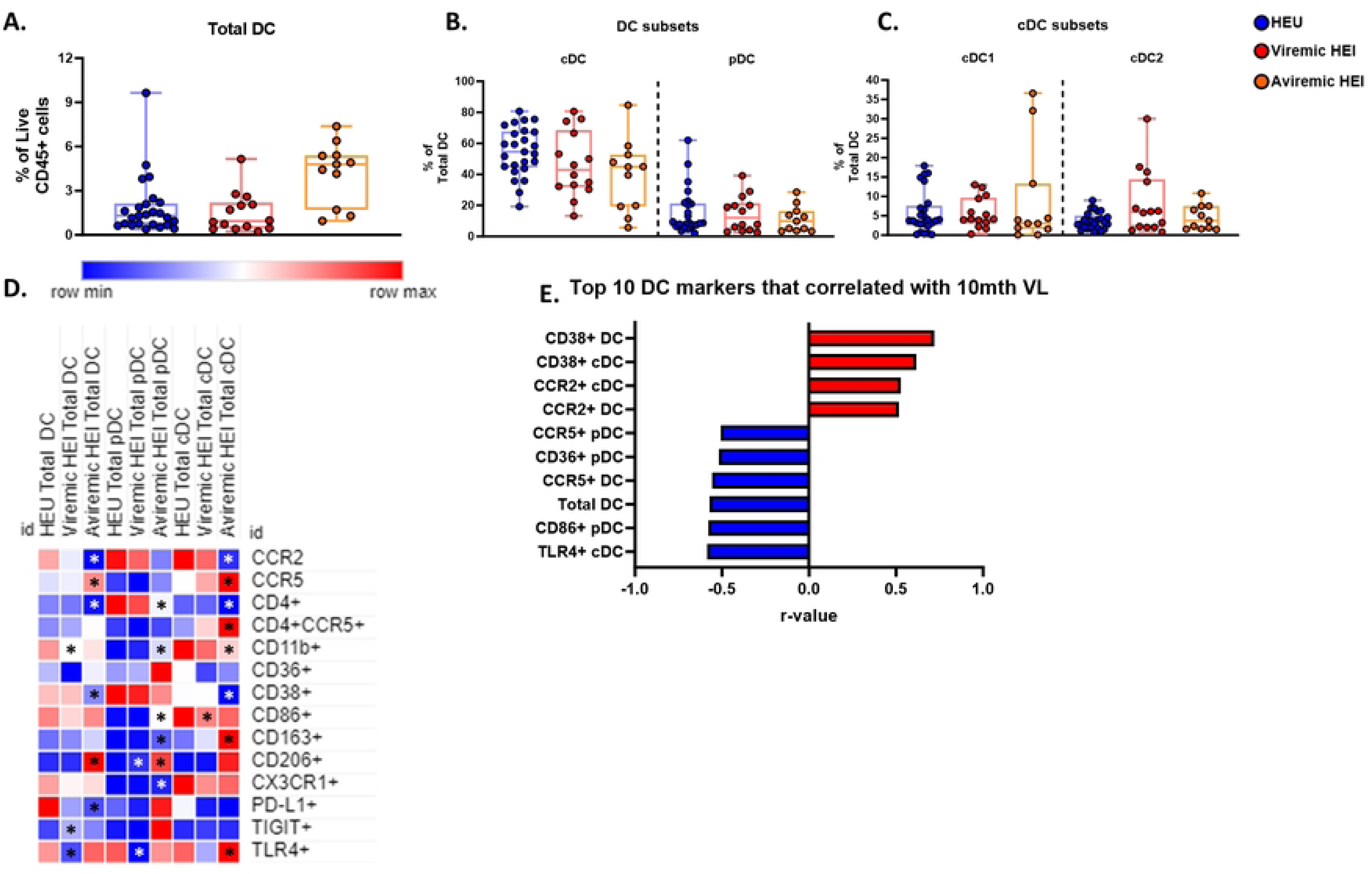
Dendritic cells of Aviremic HEI at 10 months of age show an altered phenotype compared to HEU and activation related markers correlate with plasma VL. HEI infant were split in to 2 groups at 10 months of age based on their 10-month VL. A-C) Frequency of Dendritic cells as a frequency of Live CD45+ cells, pDC and cDC as a percentage of Total Dendritic Cells, and cDC2 and cDC1 as a frequency of total cDC in HIV-exposed uninfected (HEU, blue circles, N=25), Viremic HIV-exposed infected (Viremic HEI, red circles, N=14), and Aviremic HIV-exposed infected (Aviremic HEI, red circles, N=11). D) Heatmap generated using average frequency of each marker for each respective Dendritic Cell subset in HEU and HEI without any normalization. The color gradient of the heatmap represents the average frequency of a marker between subsets with blue being the lowest frequency and red being the highest frequency. T-test Comparison were done between study groups for each marker within a row. Any significant difference for a specific marker of each subset between HEU and either Viremic or Aviremic HEI was marked with an asterisk on the heatmap. E) Top 10 most significant Dendritic Cell parameters that correlated with 10-month VL based on lowest p-value that from the Spearman Correlation and r-values of each of these correlations is plotted across the x-axis.

Interestingly, correlation of DC subsets and phenotypes with VL at 10 mo showed that all significant pDC parameters negatively correlated with VL, while cDC activation and CCR2 mediated trafficking positively correlated with VL (**Fig. 8E; Table S16**). These associations suggest that the pDC may play a role in viral control and the cDC are associated with viral replication in infants.

Longitudinal comparison of DC in 3 HEI groups (Viremic, P/L, and Early suppressed) and HEU showed cDC1 and cDC2 increase with age in HEU only and all other subsets did not change with age (**Fig. 9A-E**). Additionally, cross-sectional analysis at each timepoint revealed that frequencies of Total DC and DC subsets were similar between the 3 HEI groups and HEU (**Table S17**). At 18 months of age TLR4 expression on DC subsets was elevated in P/L group relative to other HEI and HEU participant groups but no other markers were impacted by HIV infection or virus suppression status (**Fig. 9F; Table S17**).

**Figure 9:**
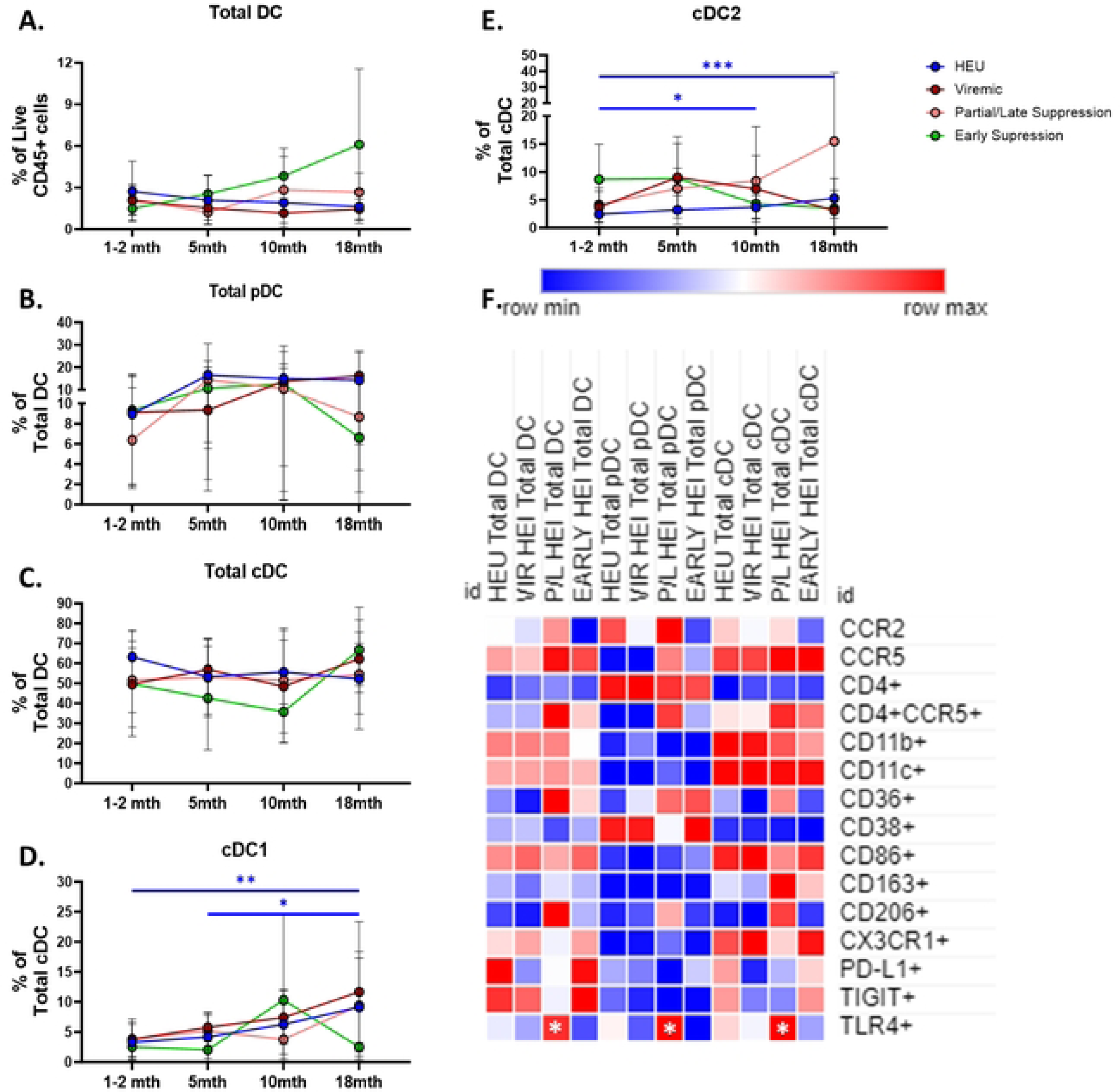
Regardless of viral trajectory, HEI show minimal differences in Dendritic Cell subset distribution and phenotype at 18 months of age compared to age matched HEU. HEI infants were split by their plasma viral load trajectories as shown in **Figure S3**. A) Frequency of Total Dendritic Cells (DC) as a percentage of Live CD45+ Cells in HEU (Blue circles, N=26), Viremic (Dark Red circles, N=13), Partial/Late Suppressors (Pink Circles, N=10) and Early Suppressors (Green circles, N=10) plotted against age in months on the x-axis. B-C) Frequency of Total pDC and cDC as a percentage Total DC in HEU, Viremic, Partial/Late Suppressors, and Early Suppressors plotted against age in months on the x-axis. D-E) Frequency of cDC1 and cDC2 as a percentage of Total cDC in HEU, Viremic, Partial/Late Suppressors, and Early Suppressors plotted against age in months on the x-axis. F) Heatmap generated using average frequency of each marker for each respective Dendritic Cell subset in HEU and HEI without any normalization. The color gradient of the heatmap represents the average frequency of a marker between subsets with blue being the lowest frequency and red being the highest frequency. T-test Comparison were done between study groups for each marker within a row. Any significant difference for a specific marker of each subset between HEU and any of the 3 HEI groups was marked with an asterisk on the heatmap.

## Discussion

Immune development during the first months and years of life is extremely dynamic, shaped by defined stages of development as well as influences of environmental and pathogenic factors. In the present study, we focused on the development of innate immune cells in infants with perinatal HIV, in consideration of the naïve status of adaptive immunity during this period. Study of innate immunity is relevant in early life because of the potential association between myeloid cells with viral persistence, and the association of NK immunity and control of HIV that previous studies have highlighted [7, 34, 35]. We identified multiple cellular markers and subsets within NK, monocyte, and DC compartments that were significantly altered in HEI and associated with viral replication. CCR2 trafficking on NK cells being negatively associated with viral replication and Intermediate Monocytes being positively associated with viral replication. Additionally, we found that their distributions and magnitude of expression of various activation, inhibitory, and trafficking markers in HEI, pre-ART initiation, were different from HEU at less than 2 months of age but became similar to HEU by 18 months of age. Our findings describe for the first time the relationship of innate immune cells and pre-ART viremia in perinatally HIV infected infants and point to a potential antiviral role for the innate immune system that could be harnessed for control of HIV in this population.

At study entry, pre-ART initiation, we found that the NK cells in HEI infants exhibited an altered profile of activation, inhibition, and trafficking receptors compared to HEU. The association of lower VL with upregulation of chemokine receptors CCR2 and CCR5, in NK cells suggests that these cells could be migrating to tissue sites with ongoing viral replication [36]. Several groups have shown that CD56^-^CD16^+^ NK cells expand during HIV infection in both children and adults [10, 37-42]. We report that infants with perinatal HIV infection also show expansion of the CD56^-^ CD16^+^ NK cells as early as 1 month of age. While the function of these cells is not fully understood, it has been demonstrated that they lack robust lytic and cytokine producing function when compared to CD56^dim^ and CD56^bright^ NK cell populations in vitro [10, 43]. Studies attempting to characterize the function of CD56^-^CD16^+^ cell subset have shown that they also produce anti-inflammatory cytokines and inhibit CD8+ T cell IFNγ production [44]. The expansion of CD56^-^ CD16^+^ NK cell subset noted in infants prior to ART at a median age of 6 weeks differs from the reduction in this cell subset observed in the very early treated Botswana infant cohort [34] who started ART within 5 days of birth and were tested at 12 weeks. These data imply that the pre-ART time point is the one which best portrays the virus host interaction.

Indeed, we showed here that the NK cells at age <2 months during ongoing viremia can kill HIV-infected cell lines. Interestingly, NK from HEI and HEU had a similar cytotoxic capacity, though the frequency of degranulating cells in response to K562 was higher in HEI. HIV reservoir analysis performed on a subset of TARA participants that achieved and maintained viral suppression with ART revealed a rapid decline in intact proviruses, relative to defective, and could be attributed to a specific vulnerability to antiviral immunity [25], possibly through NK cell activity.

Longitudinal analysis of NK cells revealed that the perturbations seen in HEI when compared to HEU at pre-ART initiation were diminished during the first 18 months of life. It has been noted in pediatric and adult studies that early initiation of ART and maintenance of viral suppression leads to at least a partial correction of persistent activation and inflammation [45-47]. a majority of the HEI in the TARA cohort had suboptimal adherence to treatment and were viremic throughout the follow up period, but our data demonstrates that many innate cellular phenotypes and subsets normalized to levels of that observed in HEU regardless of virus suppression status. In the CARMA cohort immunologic testing in adolescents who were treated within 6 months of age showed lower immune activation and better functionality compared to those who started ART after age 6 months [48]. Further investigation of the functional aspects of NK cells in the TARA cohort as well as following these individuals past 18 months of age will be needed to see how the results compare to the CARMA study.

Our results showed monocytes from HEI during pre-ART initiation showed very little alterations in the distribution of subsets and, to a lesser extent, cellular markers when compared to age matched HEU. Despite this observation, activation and trafficking markers expressed on intermediate monocytes correlated strongly with pre-ART VL suggesting a possible relationship between phenotype of monocytes early in life with viral replication, including increased frequency and MFI of CCR5 and activation marker CD38. Monocytes express markers involved with HIV entry such as CD4 and CCR5 and have been shown to be permissive to HIV infection and be a potential site of HIV reservoir. Intermediate monocytes specifically are programmed to induce a pro-inflammatory response and express higher levels of CCR5 compared to classical monocytes making them potentially more permissible to HIV infection [49, 50].

Scavenging receptors CD36 and CD206 and the integrin CD11b were also found to positively correlate with the pre-ART viral load. Studies in HIV infected adults have reported an increase in monocytes with CD36 expression compared to uninfected individuals [50-52]. In line with these reports, CM and NCM of HEI had a higher frequency of CD36 compared to the uninfected, but only IM positively correlated with VL in HEI. CD206 plays a crucial role in phagocytosis, antigen presentation, and resolution of inflammation. In our cohort CD206 was shown to positively associate with Pre-ART VL even though there was no significant difference in frequency between HEU and HEI. CD206 has been also shown in literature to be positively associated with VL in Long-Term Non-Progressors and has been negatively associated with inflammation in HIV and other diseases [53, 54]. CD11b has been shown to regulate activation and adhesion of monocytes [55]. No significant differences were observed between HEU and HEI but CD11b+ IM positively correlated with viral load at Pre-ART initiation. CD11b expression on IM were associated with impaired Ab response to flu vaccine in HIV+ adults [56]. This could potentially apply to the impairments in childhood vaccine response observed in HEI during early life [57-59]. Despite relationships of specific monocyte phenotypes with VL, we observed, over the course of the first 18 months of life, no significant differences in monocytes between HEU and HEI with minimal significant changes with age in the subsets seen in the IM and NCM suggesting the monocyte compartment is stable during the first 2 years of life.

Dendritic cells are myeloid-lineage cells important for antigen presentation and are a possible site of HIV reservoir [49, 60, 61]. This subset showed the least number of correlations with Pre-ART VL but the most alterations in the distribution of cellular subsets and markers between HEI and HEU including increased frequency and MFI of CCR5 and decreased frequency and MFI of CD11b and CD86 in HEI. Expression of CD4 and CCR5, receptors that mediate viral entry, on DC did not correlate with Pre-ART VL, but were increased in HEI and remained elevated in longitudinal analysis compared to HEU. HIV does not efficiently infect DC through CD4/CCR5 ligation but more commonly enters via DC-SIGN [29, 62-64]. DC can contribute to the viral reservoir through de novo viral production from cis-infected DC or trans-infection of CD4+ T cells through immunological synapses formed during antigen presentation or exosomes carrying infectious virions [29, 62-64]. Further studies are warranted in pediatric populations to confirm the role of DC in HIV reservoir establishment and persistence. We observed decreases in cellular markers associated with antigen presentation in DC from HEI as well which could reinforce hypothesis that responses to childhood vaccinations (e.g. Tetanus and Measles) in this population may be impaired due to impairments in APCs [57-59]. Interestingly, pDC frequencies showed a positive correlation at Pre-ART initiation, but a negative correlation with VL in our cohort at 10 months of age. It has been observed that pDC during the first 2 years of life produce lower levels of pro-inflammatory cytokines and exhibit an immature anti-viral response which could explain why a positive association was seen at study entry suggesting a potential expansion in response to infection but a negative association with VL at 10 months of age which could be possibly due to a more developed anti-viral response [7, 26, 27]. Together, these data give support towards leveraging pDC immunity in therapeutic approaches as this subset is crucial for anti-viral responses through secretion of IFN-α earlier in life when it has a more immature anti-viral response [30, 65, 66].

Our study has limitations attributable to restricted blood volumes available from infant participants, which precluded extensive multi-color flow cytometry analysis. Our overall aim was to gain a broad picture of the innate immune system; therefore we could not measure additional NK cell markers like NKG2C/D and KIR receptors, and additional activation related markers. Due to the suboptimal adherence in the TARA cohort, our study lacked robust numbers of participants with sustained virus suppression to correlate with distribution and longitudinal development of markers and subsets in HEI compared to HEU counterparts. During early life there is an increased reliance on the innate immune system to both protect from pathogens and facilitate the development of the adaptive immune system [7]. Evaluation of cellular markers and subsets during the first months of life impacted by perinatal HIV infection can provide insight into subsets most directly associated with reservoir establishment and viral control. In addition, investigating the longitudinal development of these markers and subsets could also provide insight on the potential long-term effects of HIV infection on the immune development of these infants.

## Author Contributions

SaP provided the impetus for the study through establishment of the TARA cohort with ML in Mozambique and for obtaining funding for the study. VD, LdA, SuP, SR, CD, RP, PV, ML, and SaP provided intellectual input and contributed to the experiment design. VD, LdA, and SR performed data collection. VD, LdA, AK, and SR performed data analysis and interpretation. VD, LdA, and SaP wrote the manuscript. All authors provided critical feedback to produce the final manuscript.

## Acknowledgements

The authors would like to thank all the participants in the TARA cohort and the Fundação Ariel Glaser Contra O Sida Pediátrico for their time recruiting patients and collecting samples. We thank Maria Palin and Celeste Sanchez for technical assistance in this study.

## Figure Legends

**Figure S1: Gating strategy to define NK cells and their subsets.**

**Figure S2: Gating strategy to define NK Monocytes and Dendritic Cells and their subsets.**

**Figure S3: VL trajectories of Viremic, Partial/Late Suppressors, and Early Suppressors.** HEI infants were split in to 3 groups at 18 months of age based on their VL trajectory over the first 18 months of life. A) Viremic infants were defined by never reaching a VL under 200 copies/mL during the first 18 months of life (N=13). B) Partial/Late suppression infants were defined by achieving a viral load below 200 copies/mL at a few time points throughout life or achieving and maintaining a VL below 200 copies after 12 months of age (N=10), and Early Suppressors were defined by achieving and maintaining a VL under 200 copies/mL before 12 months of age with only one rebound event (N=10). D) Table of number of participants for each group and the mean VL

